# A Python module for programmatic access to TrypTag genome-wide subcellular protein localisation data in *Trypanosoma brucei*

**DOI:** 10.64898/2026.02.17.706163

**Authors:** Ulrich Dobramysl, Richard J Wheeler

## Abstract

Protein subcellular localisation is informative for understanding potential protein function, particularly in highly structured unicellular eukaryotes. Microscopy is especially powerful for interrogating localisation, providing high resolution single cell data about where a protein resides. We previously generated the TrypTag dataset – a genome-wide protein localisation resource for the human unicellular parasite *Trypanosoma brucei* using fluorescent protein tagging. This is a puissant dataset due to its scale: Originally captured with high content image analysis in mind, it is a formidable resource for machine learning or artificial intelligence tool development and testing. Here, we describe a Python module for programmatic access to this data rich resource. Images of each tagged cell line, together with segmented cell masks, can be accessed arbitrarily by gene ID and tagging terminus, the database can be searched by protein localisation, and tools are provided to assist foundational image analysis of individual *T. brucei* cell cycle stage and morphology. We stress-tested this tool by using it to examine a key feature of *T. brucei* morphogenesis during division: The old and newly formed flagellum and associated organelles tend to have different protein compositions, and using the TrypTag toolkit we show that there is extensive age-based differential content of these organelles while the daughter nuclei completely lack such asymmetry.

## Introduction

Protein subcellular localisation is often indicative of protein function, particularly for complex and highly structured unicellular eukaryotic cells. We previously generated a genome-wide protein subcellular localisation resource, called TrypTag, for the trypanosomatid parasite *Trypanosoma brucei*^1,2^. *T. brucei* is the aetiological agent of human African trypanosomiasis (sleeping sickness) and nagana in livestock, and is one of the trypanosomatid family of human pathogens which also includes *Trypanosoma cruzi* and *Leishmania* spp. The TrypTag project was carried out as an open project for the community^3^, carried out in the procyclic form (tsetse fly gut-inhabiting) life cycle stage and used high-throughput endogenous tagging with a fluorescent protein^4^, generating cell lines expressing proteins N or C tagged from the endogenous loci. Images including hundreds of cells per tagged cell line were recorded and manually annotated, ultimately determining the subcellular localisation of 7,766 proteins – 89% of *T. brucei* protein coding genes^3^ and making a widely-used open resource^5^: TrypTag.org.

*T. brucei is* the fourth eukaryotic organism for which such a resource is available, after budding^6^ and fission^7^ yeast and human^8^ cells. TrypTag is the only such resource for a parasitic pathogen, the only such resource for a eukaryote outside of the metamonad group (which contains animals and fungi) and the only such resource for a cell with a motile cilium/eukaryotic flagellum. It was also captured with the potential for large-scale quantitative image analysis in mind, capturing more images and more cells than typical for the analogous projects in other species, and with manual expert annotation of the observed sub-cellular localisation in each cell line.

The TrypTag resource is immensely data-rich, being a database of 53,956 images covering 4,651,764 cells recording protein localisation in procyclic forms in 12,434 cell lines^1^. Most proteins were tagged at the N and C terminus, providing complementary data. We later supplemented this with a tagging of a targeted set of proteins in the bloodstream form life cycle stage, primarily with life cycle stage regulated transcripts. Although much smaller scale, we recorded similar microscopy data for 1,157 images covering 53,309 cells in 188 cell lines^9^. Microscopy of tagged proteins does bring experimental caveats: the localisation observed is that of the fluorescent fusion protein; a mutant protein. However, importantly, microscopy is a fundamentally single cell methodology in contrast to mass spectrometry-based methods for holistic subcellular localisation like LOPIT and derivatives^10^. Additionally, the per-cell data is extremely rich: a diffraction-limited fluorescent protein, fluorescent DNA stain and phase contrast image per cell. This carries enormous value for high content image analysis, especially using new artificial intelligence (AI) or machine learning approaches, with numerous recent examples for analysing cultured mammalian cells^11,11–14^. However, to succeed, the most powerful AI tools need to function across a much more diverse set of samples than only cultured mammalian cells. The TrypTag dataset also therefore poses a challenge: despite being eukaryotic cells, the *T. brucei* cell organisation is profoundly unlike the human cells in the OpenCell^15^ and Human Protein Atlas^8^ dataset.

Our previous studies have demonstrated TrypTag images as a powerful large-scale quantitative data resource to make biological discoveries. Specifically, allowing automated analysis of protein partitioning to the nucleus and nucleolus, to identify nucleolus-targeting protein properties^16^, and for semi-automated high resolution analysis of protein position within the transition fibres of the transition zone near the flagellum base^17^. Carrying out these analyses highlighted the practical challenges of working with such a large dataset. This motivated a better solution that enhances data accessibility and automation, while also emphasises the power of the dataset. We sought to solve these challenges, preparing the TrypTag dataset for future advanced analyses.

Here, we present a TrypTag Python module which handles data retrieval from the EBI BioImage Archive^18^ database (or optionally from the original TrypTag Zenodo archive^3^), allowing easy access to the localisation annotations, microscopy fields of view and individual cells for the entire dataset. Functions are provided for searching the dataset by localisation annotation and retrieving microscopy data, either fields of view or individual cells, by TriTrypDB^19^ gene ID and tagging terminus. In addition, we provide image analysis tools for measuring standard markers of *T. brucei* cell cycle stage (number of kinetoplasts and nuclei), morphology of the vermiform cell (cell length and width, kinetoplast and nucleus position) and basic fluorescent signal statistics. These can be extended with custom analyses and enable biological discoveries from the TrypTag dataset.

## Results and discussion

### Data type and structure

The raw and processed image data which went on to create the complete TrypTag dataset was originally deposited in a series of Zenodo data repositories, for each sub-dataset of a single 96 well plate. The master repository^3^ includes the table of localisation annotations and tables indexing Zenodo repositories which contain the microscopy data. Since then, the EBI BioImage Archive^18^ database became established, specifically designed for sharing large biological imaging datasets. We have deposited the processed data from the entire TrypTag procyclic form dataset^1^ (S-BIAD1866) and our targeted analysis of a smaller set of proteins in bloodstream forms^9^ (S-BIAD1932), enabling easier access to single images within these datasets (Figure 1).

**Figure 1.**
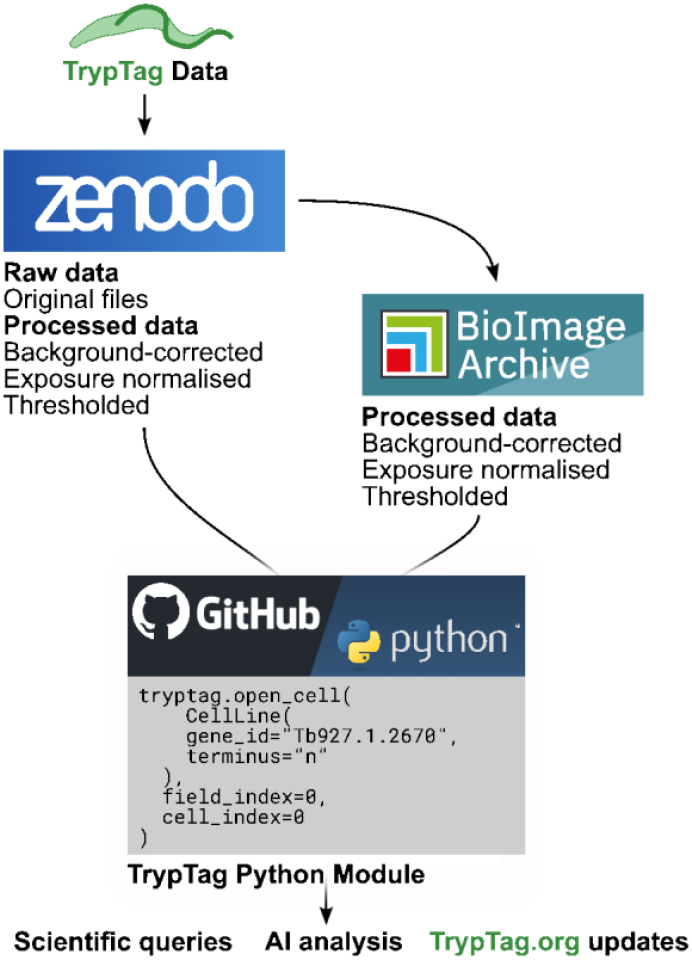
Schematic of data flow for the TrypTag Python module.

The TrypTag data is structured through 4 variables: The gene ID of the protein tagged (using TriTrypDB^19^ gene IDs), the terminus tagged (either N or C), the integer index of the field of view (typically 4 or 5 fields), and the integer index of cell within the field of view (typically tens of cells) (Figure 2A). The data is processed microscopy images, background-corrected 16-bit or 32-bit grayscale multi-slice tagged image file format (TIFF) files, with a phase contrast, tagged protein (mNeonGreen, mNG) and fluorescent DNA stain (Hoechst 33342) slice (Figure 2B). These are single focal plane images, as *T. brucei* are small and thin cells once adhered to a glass slide. The images have also been adjusted so differences in exposure time for the mNG channel have been normalised, by altering the pixel intensities proportionally. Intensity-thresholded phase contrast and DNA stain images (8-bit multi-slice TIFF files) are also provided in these repositories, as is an index table of cell locations (coordinates of a pixel within the cell mask) identified from the phase contrast image, using our previously-published conventional image analysis strategies^20^. Identification of cells in these intensity thresholded images is imperfect. Occasionally, non-cell objects may be identified as cells, distal parts of a cell may be missed (like a flagellum tip), or two touching cells may be identified as a single cell. Nonetheless, it provides practical index to a good approximation of every cell imaged for each cell line in the TrypTag dataset.

**Figure 2.**
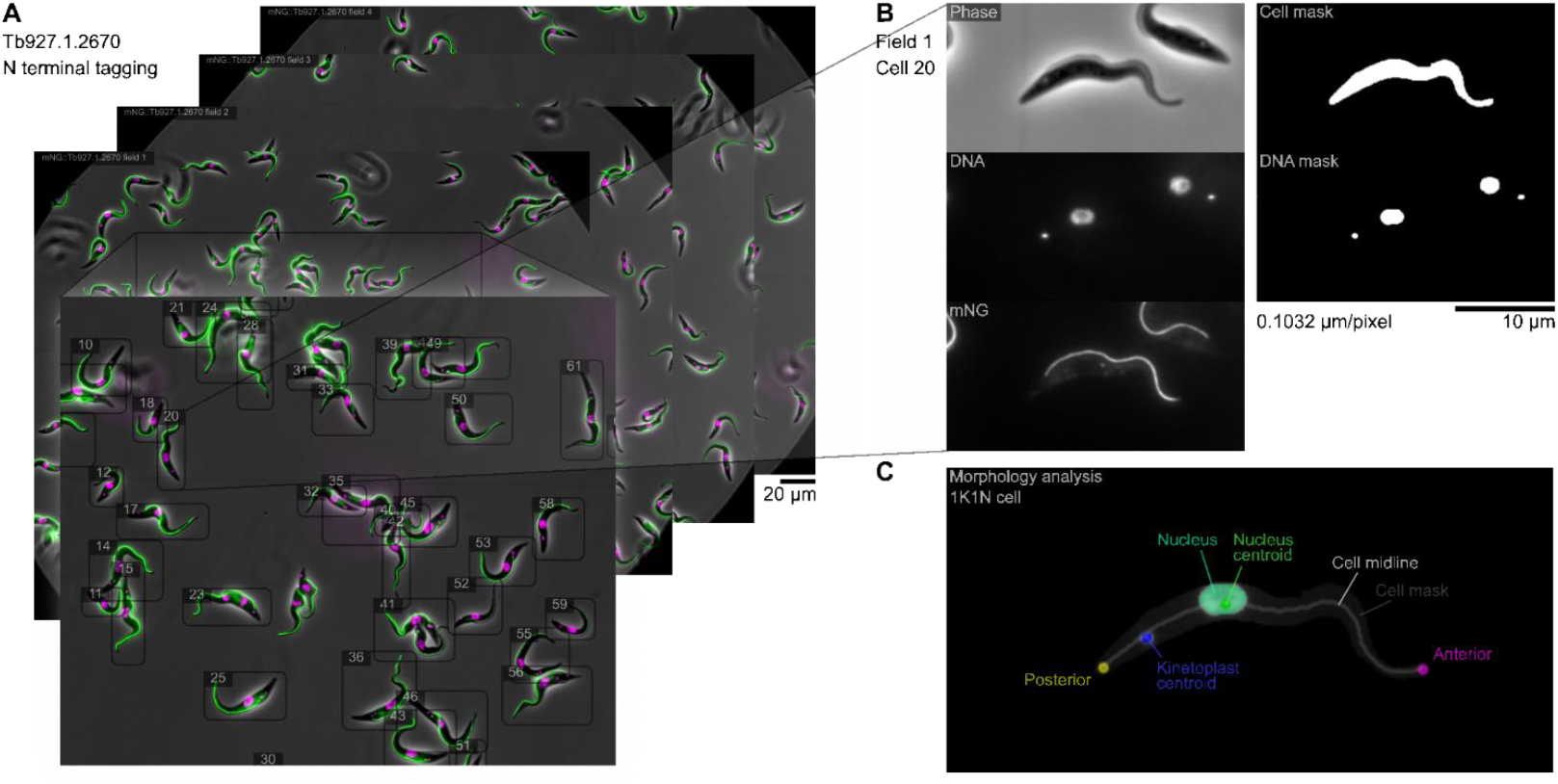
Data structure and example image data for the TrypTag dataset and Python module. **A**. Representation of the image data available for one typical cell line, indexed by gene ID (Tb927.1.2670) and by the terminus of tagging (N). This cell line was recorded with images of four fields of view, with around 50 to 80 cells per field of view. **B**. The image data available for one cell from the cell line in A. Specifically, field of view index 1 and cell index 15. **C**. Analysis of the morphology of the cell in **B** using the TrypTag Python module morphometry analysis tool, identifying the kinetoplast and nucleus, their size and position, the midline of the cell, and the positions of the kinetoplast and nucleus along it.

### The TrypTag Python module

The core purpose of the TrypTag Python module is to invisibly handle retrieval of the data from these data repositories, caching the data tables and tiff files locally, and allowing arbitrary access to images by gene ID, tagged terminus, field of view index and cell index within the field of view. Either Zenodo or BioImage Archive can be selected as the data source, with both datasets being identical. The default data source is BioImage Archive as it allows direct access to any arbitrary cell or field of view (images in the Zenodo repository can only be accessed by downloading all imaging data for a 96 well plate). The primary intended use is to analyse the TrypTag dataset, but it also supports the smaller bloodstream form tagging dataset.

The TrypTag Python module is available on Github (https://github.com/zephyris/tryptag/)^21^. Once installed and imported into a Python script, most TrypTag module functionalities are accessed by initialising a TrypTag class instance then using its methods. Here, we show some example applications, while the documentation in the Github repository (`README.md`) is the primary reference for usage. This includes instructions for installing the TrypTag module and instructions for its basic usage.

Additionally, the TrypTag Python module provides the means to carry out key analysis types. We have focused on three usage cases, which we develop with worked examples provided as iPython notebooks designed for use in Google Colaboratory. These are provided in the `examples` directory and provide a quick method for testing TrypTag module functionalities on a remote server without installing any software locally.

There are three primary functionalities, and we provide an example of each. First, using the TrypTag Python module to search for genes based on localisation annotation and accessing information about the localisation annotation term ontology (`localisation_search.ipynb`). Second, using the TrypTag Python module to open cell and field of view images for further downstream analysis in Python, i.e. using using scikit image (skimage) or numpy (`open_cell.ipynb`). Finally, pre-prepared cell image analysis tools using the integrated cell cycle stage and cell morphology analysis tools using a Python reimplementation of our previously-described morphology analysis strategies^20^ (`tryptools.ipynb`) (Figure 2C). These can be combined for high throughput proteome-wide analysis, with an example analysis provided (`analyse_list.ipynb`). Together these provide worked examples of a complete experimental analysis pipeline, from identifying cell lines by a localisation search, opening cell images from the corresponding hits, and carrying out a morphometric and signal intensity analysis of these cells.

### TrypTag.org updates

As an applied test of the TrypTag Python module functionality, we rebuilt the TrypTag.org website database. This incorporated numerous small improvements to the database, with the most important changes being updating genome annotation information to use TriTrypDB version 68^19^, adding UniProt^22^ IDs, and adding links to the EBI BioImage Archive^18^ where the high bit depth TIFFs can be downloaded. This was done in parallel with addition of the cellular component GO terms to the UniProt database^22^.

Previously, TrypTag.org website images were generated from a local copy of the TrypTag dataset using ImageJ scripts developed during the TrypTag project. We converted this to use the TrypTag Python module, accessing data from the public data repositories. The field of view images are now generated using the TrypTag Python module retrieval of field images, generating the composite images in Python. The cell cycle stage images, showing cells with different numbers of kinetoplasts (K) and nuclei (N) (1K1N, 2K1N and 2K2N cells), are now selected using the more advanced TrypTag Python module morphometry analysis. This facilitates updates, and would enable easier analysis of quantitative analyses in the future.

### Asymmetry of dividing cells

Preparation of the cell cycle stage example images re-emphasised a characteristic feature of the parasite: *T. brucei* has a highly organised cell shape, which is maintained through precise division through morphogenesis^23–28^. Specifically, in preparation for cytokinesis, the procyclic life cycle stage cell forms a new flagellum, undergoes kinetoplast (mitochondrial DNA) division, and undergoes nuclear division (mitosis). These daughter organelles are positioned along the cell: from posterior to anterior, KNKN. The kinetoplast associated with the base of the newly formed flagellum is always positioned to the posterior, and the kinetoplast associated with the base of the pre-existing (old) flagellum is always positioned closer to the middle of the cell^24^. Us and others have previously noted that there are proteins specific to the new or old flagellum and its associated structures^29,30^, potentially responsible for controlling flagellum length through the grow-and-lock model^31^. Such proteins were readily visible in the TrypTag dataset, and we noted examples beyond just the flagellar axoneme. Also numerous proteins localising to the basal body, paraflagellar rod and flagellar membrane^1^. In sharp contrast, we are not aware of any examples of proteins which are specific to one of the two daughter nuclei.

We therefore asked if, genome-wide, we could detect any proteins which are consistently preferentially present in either the more anterior or posterior nucleus of dividing cells. Using the TrypTag module, we searched for all tagged cell lines annotated as a nuclear or flagellar-associated localisation with the “cell cycle dependent” modifier. For each cell for these cell lines, we used the morphometry analysis to count the number of kinetoplasts and nuclei, selecting only cells with 2 kinetoplasts and 2 nuclei in the expected ‘KNKN’ anterior to posterior configuration for analysis. For these, we used the morphology analysis tools to identify which kinetoplast and nucleus was more anterior and which was more posterior. Finally, we measured the brightest signal within 7 μm and 11 μm of the centre of the anterior and posterior kinetoplast and nucleus respectively (Figure 3A).

**Figure 3.**
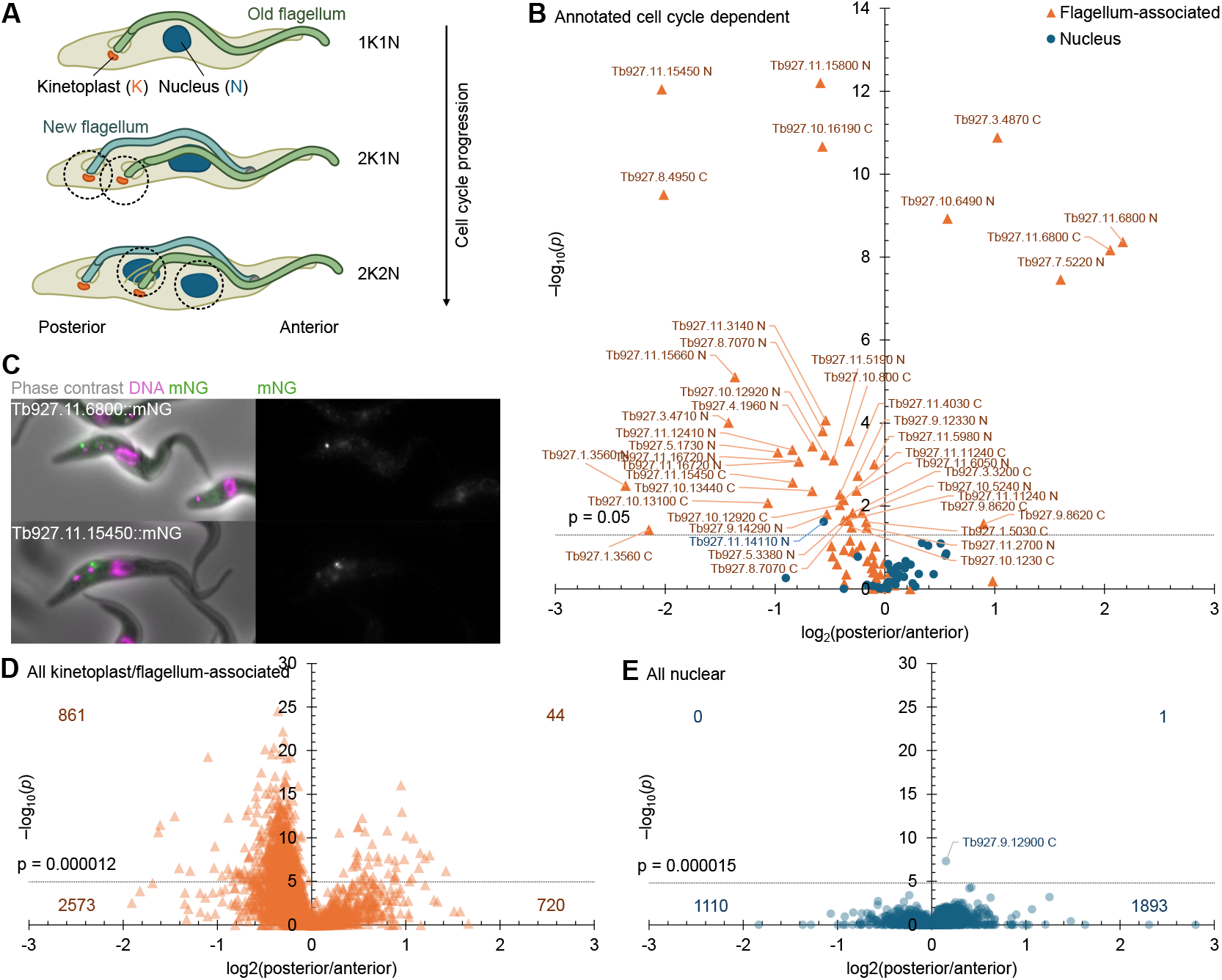
Proteome-wide detection of asymmetry in daughter organelles during cell division. **A**. Schematic of the measurement strategy to identify proteins specific to the proximal and distal kinetoplast and nucleus for flagellar-associated and nuclear proteins respectively. The posterior kinetoplast is at the base of the newly formed flagellum, the more anterior kinetoplast is at the base of the old pre-existing flagellum. **B**. Volcano plot of the ratio of maximum signal intensity near the proximal and distal kinetoplast (for flagellum-associated proteins) or nucleus (for nuclear proteins) against −log_10_ p value (two tailed T-test, no multiple comparison correction) for cell lines annotated with a localisation with a “cell cycle dependent” modifier. **C**. Example fluorescence micrographs of two cell lines with flagellum-associated proteins detected by this approach, both with basal body localisations, one new flagellum-associated and one old flagellum-associated. Left, phase contrast, DNA stain fluorescence and mNG fluorescence overlay; right, mNG fluorescence. **D**,**E**. Volcano plots of ratio of maximum signal intensity near the **(D)** proximal and distal kinetoplast (for flagellum-associated proteins) or **(E)** nucleus (for nuclear proteins) against −log_10_ p value (two tailed Mann-Whitney U test, Bonferroni multiple comparison correction) for all flagellum-associated and nuclear proteins, irrespective of “cell cycle dependent” annotations. Number of cell lines falling in each quadrant (above or below p value threshold, anterior or posterior enrichment) are indicated.

We plotted this data as a volcano plot: log ratio of posterior to anterior signal against the p value from a paired Mann-Whitney U test (Figure 3B). As selection of these cell lines was biologically motivated, we did not apply a multiple comparison correction. For flagellar-associated structures, many proteins showed a significant enrichment (p < 0.05) near the kinetoplast at the base of either the anterior (new) or posterior (old) flagellum, with more being enriched at the anterior/new. Manual inspection of these images confirmed these results, powerfully illustrated using the old and new basal body-specific proteins Tb927.11.6800 and Tb927.11.15450 (Figure 3C). In contrast, only one nuclear protein showed a (marginally) significant enrichment, and manual inspection of this suggested that it was a false positive.

To comprehensively confirm this result, we analysed every cell line annotated with a flagellar-associated or nuclear localisation, independent of whether it was manually annotated as having a cell cycle dependent localisation (Figure 3D,E, Table S1). Again, plotting log ratio of posterior to anterior signal against p value, showed numerous tagged proteins significantly enriched around the kinetoplast associated with either the old or new flagellum (Figure 3D), many more enriched at the new (861) than the old (44). In contrast only one nuclear protein was significantly enriched (Tb927.9.12900::mNG) at either the anterior or posterior nucleus (Figure 3E). The data indicate only a small enrichment (anterior 11.0% brighter than posterior), while manual inspection of the images suggest it is a false positive.

This shows that the *de novo* formation of a new flagellum and associated structures requires a large specialised protein cohort and maintenance of the old flagellum requires a smaller specialised cohort, while the division of the nucleus is a highly symmetric process, and that there do not seem to be any protein markers which demarcate one nucleus as ‘old’ and the other as ‘new’. This matches what might be expected from a binary fission of the nucleus contrasting de-novo assembly of many new flagellum-associated structures.

## Conclusions

During the development of the TrypTag Python module, we further tested it in two now published analyses: Both used it as the basis of an analysis of the distance between a flagellum-associated protein and the kinetoplast, giving insight into the ultrastructure of the base of the flagellum^32,33^.

In conclusion, the TrypTag Python module provides easy access to a large and powerful protein localisation dataset, making this resource readily accessible to future analysis like machine learning analysis of subcellular protein distribution. Our example analysis of dividing organelles shows that it can be used to ask novel biologically motivated questions about the fundamental cell biology of these parasites and readily provide quantitative answers.

## Materials and methods

All microscopy data is from the TrypTag project from *Trypanosoma brucei* TREU927 procyclic forms^3,34^. All Image analysis was carried out in Python, using scikit-image and numpy modules.

The TrypTag Python module and example iPython notebooks are available at https://github.com/zephyris/tryptag.

### Posterior and anterior kinetoplast and nucleus signal analysis

Nuclear localisations were defined as anything annotated with nucleus, or any of its child annotation terms in the hierarchical ontology^34,35^, and flagellum-associated localisations were defined as anything annotate with flagellum, kinetoplast, flagellar pocket or Golgi apparatus (or any of their child annotation terms), identified using the TrypTag Python module localisation search. For analysis of manually identified cell cycle-dependent localisation changes, only matching cell line localisation annotations with the modifier cell cycle dependent were included. Otherwise, all cell lines were included.

Morphology analysis to identify 2K1N (for flagellum-associated localisations) and 2K2N (for nuclear localisations) cells used the TrypTag Python module morphology analysis, in turn derived from our previously-described image analysis approaches^20^. This used the stringent match criteria that the morphology analysis had to identify a single unbranched midline of the cell, and that the positions of the kinetoplasts and nuclei along this midline were in the expected order from normal *T. brucei* procyclic form cell cycle morphogenesis^24^: KKN (from posterior to anterior) for 2K1N cells, KNKN for 2K2N cells. The TrypTag Python module identifies the posterior of the cell as the end of the midline closest to a kinetoplast, and subsequently the more anterior and more posterior kinetoplast (for 2K cells) and nucleus (for 2N cells) were identified based on their distance along the midline.

For signal intensity analysis, the mNG channel image was first filtered with a 7 pixel rolling ball filter (0.72 μm, all images fetched by the TrypTag Python module are at 0.1032 μm/px) to subtract both the nominal zero point of the camera (100 units intensity corresponds to 0 photons), any uniform background from scattered or out-of-focus light and any large-scale signal like uniform cytoplasmic background. The centre of the kinetoplasts and nuclei was taken as the centre of an ellipsoid fit to the masked DNA stain structure, as provided by the TrypTag Python module, and maximum signal intensity in the rolling ball background-subtracted mNG channel was measured within a 7 pixel (0.72 μm) or 11 pixel (1.13 μm) radius around the centre of the kinetoplast and nucleus.

For each cell line where at least 5 2K1N (for flagellum-associated localisations) or 2K2N (for nuclear localisations) were analysed, mean anterior mNG signal divided by mean posterior mNG signal was calculated and statistical significance determined using a Mann Whitney U test. These were plotted as log_2_ anterior/posterior signal and –log_10_ p value.

## Acknowledgements

This work was supported by the Wellcome Trust [211075/Z/18/Z, 221944/A/20/Z].

We would like to thank Nandu Gopan, Martin Weigert, Xinning Cui, Ellen Seifert and Kiran Vinayan (Technische Universität Dresden) and Ruoxuan Zeng (University of Edinburgh) for their testing of the tryptag Python module while it was in development.

## Copyright statement

This work was supported by the Wellcome Trust [Grant numbers 211075/Z/18/Z, 221944/20/Z]. For the purpose of open access, the author has applied a CC BY public copyright licence to any Author Accepted Manuscript version arising from this submission.

## Supplementary materials

**Supplemental Table 1. Data table for proteome-wide detection of asymmetry in daughter organelles during cell division**.

Data used for plotting Figure 3 D,E.

